# Differential expression of soluble receptor for advanced glycation end-products (sRAGE) in mice susceptible or resistant to chronic colitis

**DOI:** 10.1101/719310

**Authors:** Michael Bramhall, Kevin Rich, Ajanta Chakraborty, Larisa Logunova, Namshik Han, James Wilson, Catherine Booth, John Mclaughlin, Andy Brass, Sheena M. Cruickshank

## Abstract

**Aims:** Identifying the factors that contribute to chronicity in inflamed colitic tissue is not trivial. However, in mouse models of colitis, we can investigate at preclinical timepoints. We sought to validate murine *Trichuris muris* infection as a model for identification of factors that promote development of chronic colitis.

**Methods:** We compared preclinical changes in mice with a resolving immune response to *T. muris* (resistant) versus mice that fail to expel the worms and develop chronic colitis (susceptible). Findings were then validated in healthy controls and patients with suspected or confirmed IBD.

**Results:** The Receptor for Advanced Glycation End Products (*Rage*) was highly dysregulated between resistant and susceptible mice prior to the onset of any pathological signs. Increased soluble RAGE (sRAGE) in the serum and faeces of resistant mice correlated with reduced colitis scores. Mouse model findings were validated in a preliminary clinical study: faecal sRAGE was differentially expressed in patients with active IBD compared with IBD in remission, patients with IBD excluded or healthy controls.

**Conclusion:** Pre-clinical changes in mouse models can identify early pathways in the development of chronic inflammation that human studies cannot. We identified the decoy receptor sRAGE as a potential mechanism for protection against chronic inflammation in colitis in mice and humans. We propose that the RAGE pathway is clinically relevant in the onset of chronic colitis, and that further study of sRAGE in IBD may provide a novel diagnostic and therapeutic target.

## Introduction

Inflammatory bowel diseases (IBD) are a group of intestinal immune disorders, including Crohn’s disease (CD) and ulcerative colitis (UC), that cause chronic inflammation in the gut [1]. The cause of IBD is currently not known, but dysregulation of intestinal immunity, microbial dysbiosis, genetics and environmental factors contribute to disease onset. Unpredictable cycles of remission and relapse require careful monitoring and the long-term damage from inflammation often warrants potent immunomodulatory therapy or surgical intervention [2].

It is impossible to reliably predict onset, relapse or remission of IBD [3] and currently, only animal models provide a means of studying the perturbations in the gut that precede colitis. Infecting susceptible mouse strains with the enteric nematode parasite *Trichuris muris* closely parallels human Crohn’s disease in both the pathological and transcriptional changes induced [4] and has been established as a model for the study of the initiation of immune responses in the colon [5]. *T. muris* resistant BALB/c and C57BL6 mice mount an early immune response against the worms within 24 hours of infection, with large numbers of dendritic cells (DCs) migrating to the lamina propria, whereas AKR mice or C57BL/6 mice with a low dose infection mount a delayed immune response, resulting in chronic intestinal inflammation and a failure to expel the worms [5, 6]. Both susceptible and resistant strains show mild signs of inflammation within 24 hours. However, inflammation in resistant mice is controlled and ultimately resolves, whereas susceptible strains go on to develop clinical colitis in the subsequent weeks post infection.

Exploiting these early differences in the host immune response to *T. muris* infection experimentally may provide information on the factors that promote the onset of chronic, rather than resolving, inflammation in the gut [7]. Early factors are impossible to distinguish from the inflammatory milieu present in IBD patients at the point of diagnosis as chronic inflammation is already well established. Identification and validation of early changes during chronic colitis onset in mice could provide a useful pipeline for developing diagnostic and disease-management biomarkers or therapeutic targets in human colitis.

In this study, we carried out a *T. muris* infection study, investigating preclinical transcriptional changes 24 hours post infection (PI). We identified the receptor for advanced glycation end-products (*Rage*) as highly upregulated in mice susceptible to *T. muris* infection. We further investigated the presence of RAGE and related ligands in colitic mice and carried out a translational validation study investigating the presence of soluble RAGE (sRAGE) in the faeces of IBD patients and healthy controls.

## Materials and methods

### Mice

All animal procedures used in this project were carried out in accordance with the UK Animals (Scientific Procedures) Act, 1986. For *T. muris* infection experiments, 6-8 week old male BALB/c and AKR mice were used (Harlan UK, Bicester, UK). Mice were housed in individually ventilated cages with nesting material and were maintained under constant 12h light–dark cycle at 21-23 °C with free access to water and standard chow (Beekay Rat and Mouse Diet, Bantin & Kingham, Hull, UK). Euthanasia was carried out by schedule 1 procedure of CO_2_ asphyxiation followed by cervical dislocation or exsanguination. 3-6 mice were used per strain, per time point studied.

### Parasites and infection

Professor Kathryn Else, The University of Manchester, kindly provided eggs of *T. muris* Edinburgh (E) isolate for use in all infection studies. Egg infectivity and maintenance of parasite stocks were carried as described by Wakelin, 1967 [8]. Experimental mice were infected with 200 embryonated eggs in 200 μl of ultra-pure distilled water via oral gavage. Worm burden was assessed at day 21 PI. Caecum and proximal colon were harvested at autopsy to determine parasite clearance of each mouse at the end of each experiment as described by Else *et al.*, 1990 [9].

### Human samples

Prior to the commencement of the clinical study, NHS ethics approval was obtained from Berkshire B Research Ethics Committee (REC reference number: 14/SC/1413; IRAS reference number: 157778) in order to screen clinical IBD samples of faeces and serum. Participants were recruited from the Salford Royal NHS Foundation Trust. Normal healthy volunteers were recruited in accordance with the University ethics committee and the Human Tissue Act 2004. Faecal samples were taken from healthy controls (n=10) with no prior history of IBD or gut problems, or patients (n=31) with suspected IBD or clinically confirmed IBD. All patient samples were taken via outpatient clinics, returned by patients as part of standard clinical practice to be assessed for Faecal Calprotectin (FCP). Colonoscopy/biopsy were undertaken in those with elevated FCP. Of the patients, 6 patients had IBD excluded, mainly leading to a clinical diagnosis of irritable bowel syndrome (IBS), 19 known IBD patients were in remission at the of time testing (10 ulcerative colitis and 9 Crohn’s disease) and 6 patients had active IBD (n=5 CD, n=1 UC) at the time of testing.

### Statistics and analysis

Where statistics are quoted, experimental groups were compared using linear regression, Mann-Whitney U test or two-way analysis of variance (ANOVA) test followed by Sidak’s post hoc multiple comparisons test, where appropriate. P values <0.05 were considered significant. Data are presented as mean ± SEM unless otherwise stated. Statistical analyses were carried out using GraphPad Prism 7 (GraphPad Software, La Jolla, California, USA; www.graphpad.com).

## Results

### Early immune response informs resistance to *T. muris*-induced colitis

Following challenge with *Trichuris muris*, as expected BALB/c mice expelled most or all of the worms by 21 days PI, whereas AKR mice were unable to expel all worms and remained infected with a significantly higher worm burden (P=0.016, Mann-Whitney U test) (Figure 1A). Colitis scoring revealed increased histological changes associated with inflammation in both AKR and BALB/c mice after infection (Figure 1B). These changes included influx of immune cells, presence of immune cells in the submucosa, crypt hyperplasia and goblet cell loss. In agreement with previous data, the colitis scores in BALB/c mice peaked at 21 days PI and had begun to return to normal by 31 days PI. As expected, colitis scores in AKR mice rose after infection and peaked at 31 days PI where the colitis score was significantly greater than that of the BALB/c mice (P=0.046, ANOVA; Figure 1B). Representative images of haematoxylin and eosin stained proximal colon sections in naïve mice and at 31 days PI are shown in Figure 1C. Collectively, these results reproduce previously published research in the AKR/BALB/c infection model [4], where BALB/c mice initiate an acute, resolving inflammation after *T. muris* challenge and AKR mice show delayed immune response that results in a chronic inflammatory phenotype due to a failure to expel worms.

**Figure 1:**
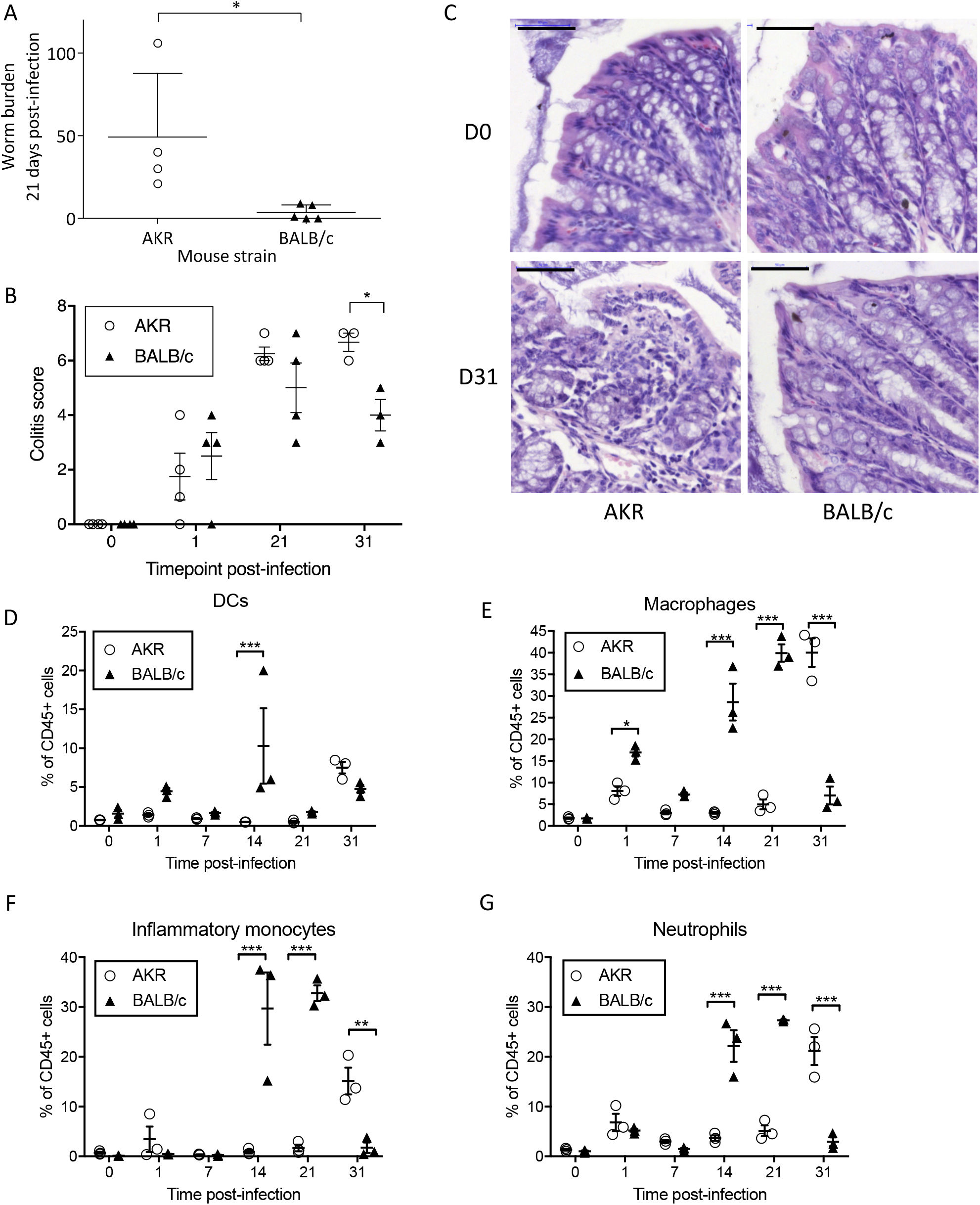
Colitis-susceptible AKR mice show delayed expulsion of *Trichuris muris* worms at 21 days and increased evidence of colitis at 31 days post infection. (A) Mean worm burden (±SD) at 21 days post infection (PI). (B) Cumulative colitis score (0-20) based on the grading of histological changes including crypt elongation (score 0-4), depletion of goblet cells (score 0-4), thickness of muscle wall (score 0-4), inflammatory cell infiltration (score 0-4) and destruction of architecture (score 0 or 3-4). (C) Representative images of haematoxylin and eosin stained proximal colon sections from naïve mice and at 31 days PI; note the high levels of immune cell infiltration and loss of goblet cells in the colonic tissues of AKR mice at 31 days post-infection. Bar=50μm. n=3-5 mice per time point. Analysis by Mann-Whitney U test or two-way ANOVA followed by Sidak’s multiple comparisons test where appropriate. (D-G) Dendritic cell (CD45^+^ MHCII^+^ CD11c^+^ F4/80^−^ CD103^+/−^ CD11b^+/−^), macrophage (CD45^+^ MHCII^+^ F4/80^+^ CD11c^+/−^), inflammatory monocyte (CD45^+^ Ly6G^+^ CD11b^+^ CD115^+^) and neutrophil (CD45^+^ Ly6G^+^ CD11b^+^ CD115^−^) populations as proportion of CD45^+^ cells (±SEM) in naïve mice and during *T. muris* challenge. n=3 mice per time point. Analysis by two-way ANOVA with Sidak’s multiple comparisons post hoc test. ***P<0.001, **P<0.01, *P<0.05.

We investigated immune cell recruitment to the colonic lamina propria by flow cytometry at 1, 7, 14, 21 and 31 days PI. BALB/c mice had an acute resolving response whereas AKR mice developed a chronic inflammatory response. At 24 hours PI BALB/c mice responded rapidly to *T. muris* challenge, with an early increase in the proportions of DCs, macrophages and neutrophils (P<0.05, ANOVA) in colonic lamina propria tissues and mesenteric lymph nodes compared to naïve mice (Figure 1D and 1E, Supplementary data). BALB/c and AKR had different responses to infection, with BALB/c tending to have greater recruitment of innate immune cells at D1 compared with susceptible AKR mice. This trend for a greater early magnitude of response in the BALB/c compared with AKR was also seen at D7, D14 and D21 post-infection with the greatest differences between the immune response of BALB/c mice to AKR at D14 PI (Figure 1B, Supplementary data). However, by D31, proportions of macrophages (P<0.001), inflammatory monocytes (P<0.01) and neutrophils (P<0.001), were all significantly greater in AKR mice, whereas the proportions of these cell types returned near to baseline levels in BALB/c mice (Figure 1D-G). The increase in immune cells observed in AKR mice at D31 corresponded with the peak colitis score, and likewise the reduction in immune cells in BALB/c mice at D31 was paralleled with a reduction in colitis score (Figure 1B). Collectively this work indicated an altered dynamic of immune response in resistant versus susceptible mice, therefore we explored the early transcriptome in order to understand changes present between colitis-susceptible and colitis-resistant mice.

### Transcriptional changes induced by *T. muris* infection identified at 24 hours post infection

Transcriptional changes in the proximal colon (the principal site of *T. muris* infection) were investigated at 24 hours post-infection via microarray, prior to the establishment of overt signs of inflammation. Genes with the largest differential expression at 24 hours post were calculated (Figure 2A). Of the 77 probe sets that were significantly upregulated in AKR mice and downregulated in BALB/c mice (1-IPPLR <0.05), 65 were successfully matched to gene IDs in DAVID. The gene upregulated in colitis-susceptible AKR mice with the largest differential expression compared to BALB/c mice was the receptor for advanced glycation end-products (*Rage*) (Log2 fold change = 2.0718; 1-IPPLR = 0.003) (Figure 2B; Supplementary data). Upregulation of *Rage* in susceptible mice was confirmed by qPCR of proximal colon at 24 hours PI in an independent experiment (P<0.001, Mann-Whitney U test, Figure 2C).

**Figure 2:**
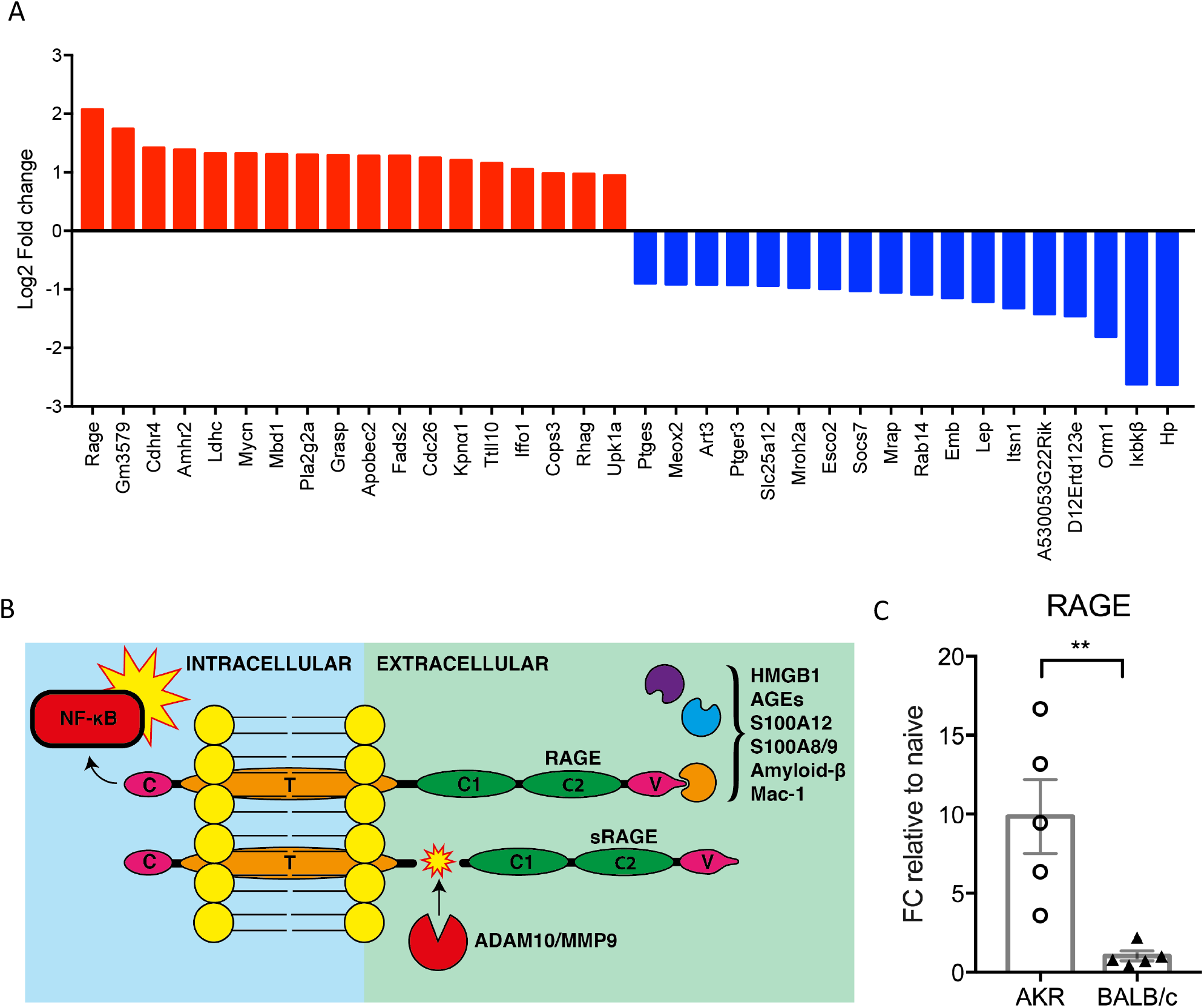
Gene expression changes in proximal colon 24 hours post-infection with *Trichuris* muris. (A) Genes most significantly upregulated in AKR mice and downregulated in BALB/c mice (red) or downregulated in AKR mice and upregulated in BALB/c mice (blue) following 24 hour *Trichuris muris* infection. (B) Schematic of the structure of RAGE, showing activating ligands, downstream NF-κB activation and formation of soluble RAGE (sRAGE) by ADAM10 or MMP9 cleavage. (C) mRNA expression of *RAGE* in the proximal colon at D1 post-infection as measured by qPCR. Data generated using Affymetrix Mouse 430 2.0 microarrays analysed using the puma and TIGERi (TFA illustrator for global explanation of regulatory interactions) packages for Bioconductor. n=4-5 mice per group. Analysis by Mann-Whitney U-test. **P<0.01.

Differential expression of transcription factors was analysed using TIGERi for MATLAB [10]. The most notable change in transcription factor gene expression was the downregulation of *FoxO4* (Forkhead box O4) following *T. muris* infection in AKR mice (Supplementary data). FOXO4 occurs downstream of the RAGE activation signalling cascade and serves to inhibit DNA binding and transcriptional activity of NF-κB (nuclear factor kappa-B), preventing inflammation [11]. *FoxO4* is also downregulated in the colonic epithelial cells of IBD patients [11]. The upregulation of proinflammatory Rage and downregulation of anti-inflammatory *FoxO4* from the RAGE signalling pathway provide compelling evidence for the relevance of RAGE activation in colitis susceptibility in AKR mice during *T. muris* infection.

### Identifying the cellular source of RAGE

Over 90% of CD326^+^ (EpCAM) epithelial cells expressed RAGE (Figure 3A) and they fell into two distinct groups, expressing either low (RAGE^lo^) or high (RAGE^hi^) levels of RAGE. In naïve mice, the total proportion of CD326^+^ epithelial cells that expressed RAGE was significantly higher in colitis-susceptible AKR mice (P<0.01, ANOVA). The proportion of RAGE^lo^ to RAGE^hi^ cells was similar in both naïve AKR and BALB/c mice, but there was significant drop in the proportion of RAGE^hi^ epithelial cells observed 2 and 7 days PI (P<0.0001, ANOVA), in both AKR and BALB/c mice (Figure 3B).

**Figure 3:**
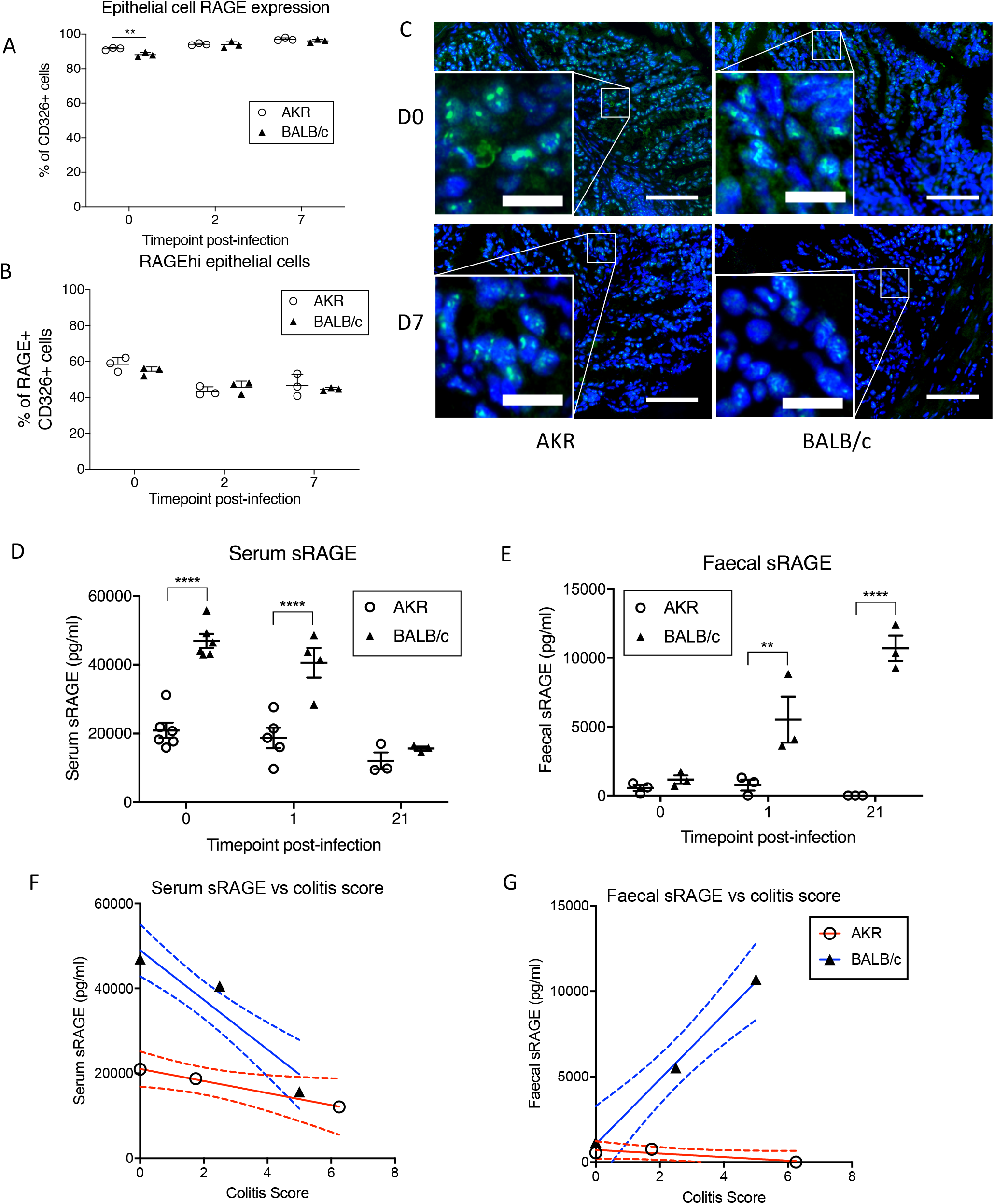
Receptor for advanced glycation end-products (RAGE) expression in the colon of *Trichuris muris* infected AKR and BALB/c mice (aged 6-8 weeks). (A) Proportion of RAGE expressing epithelial cells (CD326^+^, ±SEM) during early *Trichuris muris* infection. (B) Proportion of colonic epithelial cells expressing high levels of RAGE is reduced shortly after infection, as measured by flow cytometry. (C) Representative images of colon sections stained for RAGE (FITC; green) and nuclei (DAPI; blue) in naïve mice and at 7 days post-infection (Bar = 100μm; inset bar = 22μm). (D-E) sRAGE present in serum or faeces during *Trichuris muris* infection as measured by ELISA. (F-G) Correlation of serum and faecal sRAGE versus colitis score at 0, 1 and 21 days post-infection. n=3-5 mice per time point. Analysis by linear regression, two-way ANOVA with Sidak’s post hoc test. **P<0.01 ****P<0.001.

Immunohistochemistry was used to confirm expression of RAGE in tissue (Figure 3C). Akin to the flow cytometry data, we saw high expression of RAGE throughout the colonic epithelium. Concurring with the flow cytometry data, there was only minimal fluorescence seen in the immune lamina propria cells indicating that epithelial cells indeed express greater amounts of membrane-bound RAGE than immune cells. This data suggest that the epithelial cells are likely to be the source of the observed increase in *Rage* mRNA. There was a reduction in intensity of RAGE staining at 7 days PI by immunohistochemistry (Figure 3C) compared to naïve mice, which also correlated with the measured shift in proportions of epithelial cells from RAGE^hi^ to RAGE^lo^ cells measured by flow cytometry.

### Is RAGE differentially cleaved in colitis-susceptible and colitis-resistant mice?

RAGE may be internalised after ligand binding, or released as soluble RAGE (sRAGE) via enzymatic cleavage by ADAM10 or MMP9 [12, 13]. To investigate whether RAGE was being cleaved we assessed sRAGE levels in serum and the faeces by ELISA. Serum sRAGE levels in susceptible mice remained constant throughout the experiment. Resistant mice had significantly higher serum sRAGE than susceptible mice both prior to infection and at 24 hours PI (P<0.001, ANOVA; Figure 3D). sRAGE was also detectable in faeces, where both resistant and susceptible mice had very low levels of sRAGE prior to infection. Over the course of infection, faecal sRAGE in BALB/c mice increased to significantly higher levels than susceptible mice at 24 hours and 21 days PI (P<0.01, ANOVA; Figure 3E). Faecal sRAGE was not detected at all up to D21 in susceptible mice.

Correlation of serum and faecal sRAGE levels to colitis scores highlights the changing levels of sRAGE in BALB/c mice during the course of *T. muris* infection relative to pathological changes in the colon (Figure 3F-G). The increased circulating serum sRAGE at D0 in BALB/c mice, where colitis scores are lowest, drops as colitis increases at D1 and D21 (R^2^=0.90). Faecal sRAGE in the BALB/c mice increases relative to colitis scores (R^2^=0.99). However, sRAGE levels in susceptible mice did not change during the course of infection relative to increasing colitis scores from D0 to D21 post-infection (serum R^2^=0.99, faecal R^2^=0.73).

We then investigated levels of the RAGE ligand S100A8 (one part of the heterodimeric calprotectin protein, currently used as a clinical biomarker for IBD) in serum and faeces as an indicator of whether sRAGE might be quenching the proinflammatory effects of circulating RAGE ligands by acting as a decoy receptor. Serum and faecal S100A8 did increase slightly during the course of infection in both AKR and BALB/c mice, but no statistical differences were observed between the two strains and there was high variability between mice. At 21 days PI BALB/c mice had greater levels of serum S100A8 than AKR mice (not significant, ANOVA; Supplementary data). Faecal S100A8 remained broadly similar in naïve and *T. muris* infected mice of both AKR and BALB/c strains. As with serum S100A8, faecal S100A8 was slightly raised in BALB/c mice at 21 days PI compared to AKR mice but this was not significant (ANOVA; (Supplementary data)). In addition to being highly variable, S100A8 correlated poorly with colitis scores (Supplementary data).

### sRAGE is differentially expressed in IBD

As the differences in sRAGE were most apparent and consistent in faeces we focused on analysis of faecal specimens from healthy volunteers and patients with IBD or suspected IBD. sRAGE was not detected in the faecal samples of healthy volunteers. In contrast, s-RAGE was detectable in the patient cohort (Figure 4A). The highest levels of sRAGE were seen in patients with IBD excluded (largely IBS ascribed) and IBD in remission, although remission patients were more variable. Patients with active IBD characterised by severe inflammation had low levels of sRAGE and increased calprotectin.

**Figure 4:**
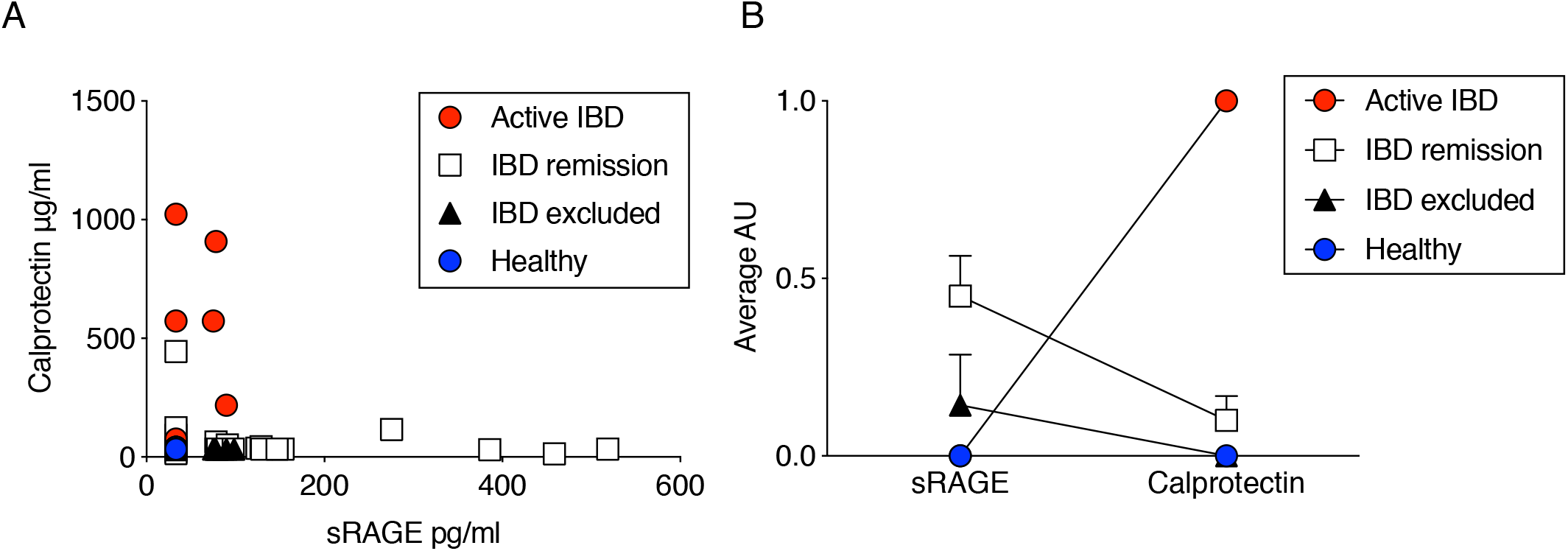
Soluble receptor for advanced glycation end-products (sRAGE) is detectable in the faeces and serum of IBD patients. (A) Scatter plot of sRAGE (pg/ml) versus calprotectin (μg/ml) present in the faeces of patients with active IBD, IBD in remission, IBD excluded (IBS) compared to healthy controls. (B) Relative levels of faecal sRAGE versus calprotectin in human IBD/IBS or healthy controls. Data were transformed to arbitrary units (AU) where samples greater than 3SD from baseline = 1 (present), otherwise they scored 0 (absent).

We then transformed the faecal RAGE and calprotectin ELISA data into present (1) or absent (0), where protein is scored as present if ≥ 3*SD above baseline. When patient data was stratified into groups (active IBD, IBD in remission, IBD excluded (IBS) and healthy controls) the ratio of RAGE to calprotectin clearly identified healthy controls (0, 0) and active IBD (0, 1) from IBS and IBD in remission (Figure 4B). In healthy individuals, we saw no sRAGE or calprotectin signal in any subject. In active IBD we had a consistent pattern of Calprotectin present, but no sRAGE. For patients whose disease is resolving we saw a more complicated picture, where we had a subset of patients undergoing routine test that were expressing both sRAGE and Calprotectin.

## Discussion

Animal models are crucial for examining the causative events that lead to diseases such as chronic colitis. Genetic factors influence the likelihood of developing colitis, but it is impossible to continually monitor individuals with potential genetically susceptibility in order to identify the pathways that drive the development of chronic intestinal inflammation. Models that accurately simulate preclinical changes in the gut allow us to interrogate the pathways leading to chronic colitis, prior to the development of the complex inflammatory environment when disease is established. Levison *et al.* [14] have previously shown that *T. muris* infection in AKR mice causes colitis that correlates phenotypically and transcriptionally with the profile of human CD. Here, we provide novel evidence reinforcing the use of the *T. muris* infection as a model for the discovery of preclinical intestinal inflammation markers that translate into human IBD patients.

In line with previous *T. muris* infection studies, we observed altered dynamics of the immune response between colitis resistant versus susceptible mice, characterised by an early influx of DCs [5]. We then identified upregulation of *Rage* as a potential indicator of colitis susceptibility in mice. RAGE activity has already been linked to active IBD [15] as well as other inflammatory diseases including diabetes, Alzheimer’s, airway inflammation, cancer and haemorrhagic shock [16]. Additionally, several RAGE ligands have been identified as associated with the inflammation in IBD, including calprotectin, EN-RAGE and HMGB1 [17–19]. Thus, our data from the mouse model shows a clear link to known pathology in IBD. It is important to validate results from mouse model to human disease and therefore we conducted a small validation study to assess faecal sRAGE. Faecal sRAGE was readily detected in patient’s faecal samples. Akin to the mouse data we saw lower sRAGE in patients with active chronic inflammation. Surprisingly, symptomatic patients with IBD excluded on a basis of normal FCP and/or colonoscopy had higher levels of sRAGE than healthy volunteers. Largely, IBS was clinically ascribed but that was not based on formal diagnostic criteria, and other diagnoses such as bile acid diarrhoea or microscopic colitis were not formally excluded. The sample size was small and more prospective studies are needed to confirm this preliminary observation. Patients whose IBD was reported to be in remission had variable levels of sRAGE. It is tempting to speculate that lower levels of sRAGE are associated with a risk of subsequent flare of inflammation but as we only had single samples from the patients, we cannot assess this, however it would be interesting to track sRAGE over time in a prospective study of patients with IBD.

RAGE has been described on several immune cells with, for example, neutrophils identified as expressing large amounts of RAGE [20]. Our flow cytometry and immunohistochemistry analysis of RAGE expression in multiple cell types present in and around the lamina propria and crypts of the colon, however, did not suggest that immune cells were the main sources of cellular RAGE. Our data in fact showed that a major cellular source of RAGE in the gut were the gut epithelial cells. Previous studies have shown that epithelial cells not only express RAGE, but also upregulate RAGE expression during colonic inflammation [15]. Changes we observed in the levels of RAGE expression suggest that it is the epithelial cell response to *T. muris* infection that informs subsequent susceptibility to chronic inflammation. The reduction in the amount of RAGE present at the cell membrane we saw by flow cytometry and by immunohistochemistry immediately following *T. muris* infection could be caused by either internalisation of activated RAGE-ligand complexes or ADAM10-mediated shedding to produce sRAGE [21, 22]. Splice variants of *Rage* may also result in a truncated RAGE molecule or a modified and actively secreted decoy receptor [23, 24]. The process by which epithelial cells may undergo RAGE shedding represents an important distinction in the course of gut immunity and homeostasis, and may be an essential component in dictating whether inflammation becomes chronic or resolves. Further investigation into the extent to which splice variants, internalisation or sheddase activity form the mechanism for changes in sRAGE in both the *Trichuris* model and in human patients could provide further insight for the role of the RAGE pathway in colitis.

Activation of RAGE can have several outcomes including immune cell migration and transcription of pro-inflammatory cytokines. RAGE-mediated leukocyte migration via CD11b (Mac-1) is involved in migration of immune cells to the site of injury and homing of DCs to the lymph nodes [15, 25]. However, whether the upregulation of *Rage* is crucial to facilitate the early DC migration in the *T. muris* model remains unclear. While we observed differences in DC migration as early as day 1 PI, we saw no differences between cell surface RAGE expression between AKR and BALB/c mice that might account for the altered dynamics of DC recruitment. Similarly, RAGE has been linked to neutrophil recruitment but we only observed modest neutrophil infiltration in the first 24 hours PI and no difference between colitis-susceptible or colitis-resistant mice. Despite minimal differences in immune cell presence during the early stages of *T. muris* infection, prolonged activation of RAGE results in activation of inflammatory signalling molecules including NF-κB and MAP kinases [16]. Consequently, an environment where RAGE ligands such as HMGB1, and S100 proteins are continually present results in perpetual NF-κB activation and subsequent chronic inflammatory conditions.

We observed striking differences in the levels of faecal and systemic sRAGE between colitis-resistant and susceptible mice, with BALB/c mice rapidly producing sRAGE in response to *T. muris* infection. sRAGE effectively acts as a decoy receptor for RAGE as it can still bind to the same damage induced ligands as membrane bound RAGE but, as it lacks a cytoplasmic tail, it cannot initiate the pro-inflammatory signalling cascade [26]. Thus, higher levels of sRAGE might be expected to block inflammation and indeed reduced levels of sRAGE have been found in mice with chronic inflammation as well as patients suffering from chronic inflammatory diseases [27]. The reduction in epithelial RAGE expression followed by increases in circulating sRAGE in colitis-resistant mice suggests shedding of RAGE to form sRAGE as a protective feedback process against the development of chronic inflammation. Colitis-susceptible AKR mice show the same reduction in epithelial RAGE expression but did not produce sRAGE in the same quantities, suggesting internalisation and activation of RAGE and initiation of subsequent proinflammatory pathways after *T. muris* infection. Indeed, it is known that *T. muris* excretory/secretory (E/S) products induce NF-κB signalling in colonic epithelial cells shortly after infection and the susceptible immune response to *T. muris* is associated with expression of the T helper (T_H_) 1 cytokines interferon (IFN)-γ, tumour necrosis factor (TNF)-α and IL-12 [28]. This response parallels the cytokines expressed after RAGE activation, which include T_H_1 and T_H_17 cytokines TNF-α, IL-1α, IL-6, IL-8 and IL-12 [16]. Diagnosis of IBD usually involves an assessment of clinical history and physical examination, with endoscopy and histology considered to be the gold standard tools [29]. Accurately assessing disease activity remains dependent on colonoscopy and/or small bowel imaging. The invasiveness of current diagnostic methods is not ideal and recent work has aimed to identify serum or faecal biomarkers that can reliably identify active disease [30]. The number of potential IBD biomarkers is high, but there remains a lack of reliable and reproducible biomarkers for use in clinical practice [31]. Efficacy of current therapies is also variable, with risks of sometimes serious side effects, especially infection meaning there is considerable interest for new biomarkers and new therapeutics [32]. Calprotectin entered clinical practice as an IBD biomarker to aid clinical diagnosis non-invasively, but measurements of faecal calprotectin are variable and there is little agreement about what should be considered a normal baseline level in healthy patients [33]. Indeed, the concept of a simple normal cut off is impossible to entertain given the enormous heterogeneity in faecal water content, matrix composition, transit time, site and extent of inflammation and the contact of faecal component sampled with the mucosa; composite measures are essential. Calprotectin is a product of tissue damage and binds to RAGE to promote inflammation, but this action will be reduced in the presence of the decoy receptor sRAGE. Our preliminary observation of alterations in sRAGE expression in mouse and human disease suggests there may be merit in looking at both calprotectin and sRAGE to better predict whether calprotectin and other RAGE ligands are indeed able to drive pro-inflammatory signals. This may improve the reliability of calprotectin as a biomarker.

Our pilot clinical study successfully validated the use of the *T. muris* infection model as highly translatable to human IBD states. The *T. muris* model is a useful tool in dissecting early pathways that are involved in the onset of colitis. By using a mouse model and focusing on early initiating events in the development of colitis, we have identified a potential role for RAGE in mediating the development of inflammation. Furthermore, our observation of high levels of sRAGE in acute resolving inflammation suggested there may be utility in monitoring of sRAGE to monitor IBD. However, a larger clinical study would be required to investigate this further.

## Acknowledgements

Funding for this research was provided by the EPSRC, Epistem Ltd. and the BBSRC impact accelerator fund. NHS Salford Trust provided the human clinical samples for this project, and we are grateful to all clinical staff, patients and healthy volunteers who took part in the trial.

The Bioimaging Facility microscopes used in this study were purchased with grants from BBSRC, Wellcome and the University of Manchester Strategic Fund. Special thanks go to Roger Meadows from the Bioimaging Facility for his help with the microscopy and image acquisition.

