## Supplemental information for "Differential expression of soluble receptor for advanced glycation end-products (sRAGE) in mice susceptible or resistant to chronic colitis"

**Supplementary Information**

**Supplementary materials and methods**

**Immunohistochemistry**

**Histology**

Proximal colon snips were fixed in neutral buffered formalin (NBF; 10% neutral buffered formalin in PBS) for 24 hours, processed (Shandon Citadel 2000; ThermoShandon, Runcorn, UK) and embedded in paraffin wax. 5 μm sections were then dewaxed, rehydrated and stained using a standard Haematoxylin & Eosin (H&E) stain or Alcian blue/Periodic Acid Schiff’s stain [1]. Images were acquired using a 20x/0.80 Plan Apo objective using a slide scanner (Pannoramic 250 Flash II; 3D Histech, Budapest, Hungary) and samples were measured for crypt hyperplasia, immune cell infiltration, immune cell location, goblet cell loss and muscle wall thickening using Pannoramic Viewer software (3D Histech). All slides were measured and counted blind and in a randomised order. Colitis scoring was carried out following a methodology described previously [2]. Briefly, sections received a combined score for crypt elongation (score 0-4), depletion of goblet cells (score 0-4), thickness of muscle wall (score 0-4), inflammatory cell infiltration (score 0-4) and destruction of architecture (score 0 or 3-4). Normal, healthy colon would score 0-1, and mild inflammation would score 5-6 (Supplementary Table 1).

**Immunofluorescence**

Antibodies specific to RAGE (Abcam, Cambridge, UK), ADAM10 (R&D Systems Europe, Abingdon, UK) and cytokeratin (Sigma-Aldrich, Poole, UK) were used to detect RAGE expression, ADAM10 protein expression and epithelial cells. 5 μm sections of proximal colon were fixed in 4% paraformaldehyde and blocked using the tyramide blocking kit (Perkin Elmer, Cambridge, UK), incubated with primary antibodies for 60 minutes, and FITC labelled secondary antibodies for 60 minutes and mounted using Vectashield pro-long anti-fade reagent containing the nuclear counter-stain 4′,6′-diamidino-2-phenylindole (DAPI; Vector Laboratories).

Fluorescence imaging was carried out using an Olympus BX51 microscope using either a 10x/0.30 UPlanFLN objective or a 40x/0.75 UPlanFLN objective with DAPI, FITC and Cy3 filters, coupled with a CoolSNAP EZ camera (Photometrics, Tucson, USA). Specific band pass filter sets for DAPI, FITC and Cy3 were used to prevent bleed through from one channel to the next. Images were first captured on MetaVue (Molecular Devices, Sunnyvale, USA) and then corrected using ImageJ64 v1.44o (National Institute for Health) with ‘ImageJ for Microscopy’ plugins (McMaster Biophotonics Facility, Hamilton, Canada).

**Large intestine cell isolation and flow cytometry**

Caecum and colon were digested in RPMI-1640 containing 5% L-glutamine, 5% penicillin/streptomycin, 10% foetal bovine serum (Sigma-Aldrich), collagenase (1 mg/ml; type VIII from *Clostridium histolyticum*; Sigma-Aldrich) and dispase (0.5 mg/ml; Gibco, Paisley, UK). Cells were filtered and separated by density gradient centrifugation as described by Bowcutt *et al.* 2014 [1]. Resultant cells were resuspended in FACS buffer (PBS containing 0.5% BSA and 0.1% NaN_3_; Sigma-Aldrich) prior to staining and flow cytometry acquisition. MLN cells were dissected and filtered and resuspended in FACS buffer prior to staining and flow cytometry acquisition. Fc receptors were blocked using 2 μg/ml anti-CD16/32 (eBioscience, Hatfield, UK) and immune cells stained with antibodies specific to CD115, CD11b, CD45, Ly6G, RAGE, CD11c, F4/80, MHCII, CD103 and CD326 (EpCAM). Cells were acquired by flow cytometry using an LSRII (Becton Dickinson). Data was analysed using FlowJo v10 flow cytometry software (Tree Star, Oregon, USA).

**Faecal protein extraction and preparation**

Protein from faecal samples were extracted using stool prep tubes (Cat: BS-00-03; Bioserv, Rostock, Germany) according to the manufacturer’s protocol. 100 mg of faecal sample was treated with extraction buffer and resultant faecal suspensions were spun at 15000 rcf for 5 minutes. Supernatant was then transferred to Amicon Ultra-2 Centrifugal Filter Units with Ultracel-10 membranes (Cat: UFC201024; Millipore, Watford, UK) and spun at 2800 rcf for 15-20 minutes until 100 μl of concentrated sample remained.

**Serum preparation**

Approximately 1 ml of blood was drawn and left to clot at room temperature for 1 hour. Samples were spun at 15000 rcf for 10 minutes and serum stored at -20^o^C.

**RAGE and S100A8 ELISA**

Faecal and serum ELISAs were carried out to measure RAGE (Cat: DY1179, R&D Systems, Abingdon, UK) and S100A8 (Cat: DY3059, R&D Systems) according to the manufacturer’s protocol. Briefly, 96-well ELISA plates were coated with capture antibody (RAGE: 1.0 μg/ml in 0.22 μm filtered PBS; S100A8: 4.0 μg/ml in 0.22 μm filtered PBS), incubated overnight, washed and blocked with reagent diluent (1% BSA in PBS, pH 7.2-7.4, 0.22 μm filtered before incubation with duplicate faecal extracts and serum samples. A seven-point standard curve was prepared according to recommendation and negative controls of reagent diluent only were used. Samples were incubated for 2 hours at room temperature, washed, and detection antibody added for 2 hours. Plates were washed, and visualised using Streptavidin-HRP (Cat: DY998, R&D Systems) and substrate solution and stop solution (2 N H_2_SO_4_). The optical density of each well was read immediately, using a microplate reader (MRX II, Dynex Technologies, Worthing, UK) set to 450nm. Readings at 570nm were subtracted to correct for optical imperfections in the plate, as per the manufacturer’s instructions.

**RNA extraction for PCR**

100 mg colonic tissue samples were snap-frozen in 1 ml TRIsure (Cat: BIO-38032, Bioline, London, UK) and stored at -80^o^C until use. Samples were transferred to lysing matrix tubes (MP Biomedicals, Fisher Scientific, Loughborough, UK) containing 1.4mm ceramic spheres and homogenised twice using a FastPrep-24 (MP Biomedicals, Fisher Scientific, Loughborough, UK) before adding 200 μl of chloroform (Sigma-Aldrich), mixing vigorously and spinning for 15 minutes at 12000 x *g*. RNA in the upper aqueous phase was precipitated by adding chilled (4^o^C) isopropyl alcohol. After incubation and centrifugation, RNA was washed with 75% ethanol and resuspended in 25 μl of nuclease-free water (NFW; Life Technologies, Paisley, UK). RNA quality check and yield was measured (Nanodrop ND-1000, Thermo Scientific, Wilmington, USA). Prior to cDNA conversion, genomic DNA was removed by incubating with RQ1 RNase-free DNase (Cat: M610A, Promega, Southampton, UK) at 37^o^C for 30 minutes. 2 μl DNase stop solution (Promega, Southampton, UK) was added and samples were incubated at 65^o^C for 10 minutes and suspended in NFW.

**cDNA conversion**

RNA was converted to cDNA using a BioScript reverse transcriptase kit (Bioline, London, UK) according to the manufacturer’s recommendations. Samples were run on a Peltier thermal cycler (MJ Research PTC-225, Cambridge, USA) on the following cycle: 25^o^C for 5 minutes, 40^o^C for 60 minutes, 70^o^C for 15 minutes and stored at -20^o^C until quantification via qPCR. Endpoint PCR and optimisation was carried out using a MyTaq HS DNA polymerase kit (Bioline, London, UK) according to the manufacturer’s protocol.

**qPCR**

qPCR was conducted using a KAPA SYBR Fast qPCR kit (Cat: KM4105, Kapa Biosystems, Wilmington, USA) as per the manufacturer’s protocol. Primer pairs used were RAGE forward: 5’-CCAATGGTTCCCTCCTCCTT-3’; RAGE reverse: 5’-TTGGACTTGACCTCCTTCCC-3’ (Eurofins, Ebersberg, Germany). Plates were run on a single-colour real-time PCR detection machine (MyiQ, Cat: 170-9740, BioRad, Hemel Hempstead, UK) at 95^o^C for an initial denaturation cycle for 5 minutes, followed by 35 cycles of 95^o^C denaturation for 15 seconds, 59^o^C annealing for 15 seconds and 72^o^C extension for 10 seconds.

**RNA extraction and quality analysis for microarray**

Five mm sections of proximal colon were stored in 500 μl RNALater (Invitrogen, Paisley, UK) at 4^o^C overnight and then transferred to -80^o^C. Samples were then defrosted and homogenised using a bead mill. RNA was extracted using an Ambion RNAqueous Kit (DNase treated) according to the manufacturer’s instructions. RNA quality check for integrity and yield was performed using a NanoDrop spectrophotometer (Labtech International, East Sussex, UK), Agilent Bioanalyser and MultiNA (Shimazdu, Milton Keynes, UK). An RNA integrity number (RIN) of 7 was used as a cut-off for RNA quality; all samples with an RIN of ≥7 progressed to cDNA conversion using Epistem RNAamp reverse transcription amplification.

**Microarray hybridisation**

Gene expression was determined using a 45101 probe-set murine oligonucleotide array (Affymetrix mouse 430 2.0 microarray). Hybridisation of samples to arrays and array scanning was carried out by Epistem Ltd according to the manufacturer’s instructions. Samples were not pooled; each array was produced using cDNA from a single mouse.

**Microarray analysis**

All microarray analyses, including pre-processing, normalisation and statistical analysis were carried out using R [3] version 3.2.1 and Bioconductor [4] version 3.1. Data quality was assessed using AffyQCReport [5] and microarray data were pre-processed using the multi-chip modified gamma Model for Oligonucleotide Signal (mmgMOS) normalisation method (with all options set to default). Differential expression was analysed using pumaComb and pumaDE, both implemented within the Propagating Uncertainty in Microarray Analysis (puma) [6] Bioconductor package. Outputs contained log2 fold change values and improved probability of positive log ratios (IPPLR); statistical significance values generated in the puma package. Resulting gene lists were converted from Affymetrix probeset IDs to official gene symbols using NetAffx [7] and submitted to the Database for Annotation, Visualization and integrated discovery (DAVID) [8, 9] and mapped to KEGG biological pathways [10, 11].

Analysis of differential expression of transcription factors was carried out using the TIGERi (TFA illustrator for global explanation of regulatory interactions) [12] package for MATLAB [13] using the default settings.

**Supplementary Table 1:** Histological scoring system for mucosal inflammation.

| Crypt elongation | Goblet cell depletion | Muscle wall thickness | Inflammatory cell infiltration | Destruction of architecture |
| --- | --- | --- | --- | --- |
| 0: Normal crypt length | 0: Normal goblet cell numbers | 0: Normal thickness | 0: No or occasional inflammatory cells in the lamina propria | 0: Normal architecture |
| 1: Up to 25% increase | 1: Up to 25% decrease | 1: Up to 25% increase | 1: Moderate number of inflammatory cells in the lamina propria | - |
| 2: 25%-50% increase | 2: 25%-50% decrease | 2: 25%-50% increase | 2: High numbers of inflammatory cells in the lamina propria | - |
| 3: 50%-75% increase | 3: 50%-75% decrease | 3: 50%-75% increase | 3: Confluent inflammatory cells in the lamina propria | 3: Crypt deformity |
| 4: >75% increase | 4: >75% decrease | 4: >75% increase | 4: Confluent Inflammatory cells in the lamina propria extending to the submucosa | 4: Gross structural changes including ulcers |

**Supplementary Table 2.** Human control and patient details

| **Diagnosis** | **n** | **Sex** | **Age** | **Disease history** |
| --- | --- | --- | --- | --- |
| Healthy control | 10 | 7M, 3F | 20-56 | None |
| IBD excluded | 6 | 2F, 4 not reported | 30-39 | IBD excluded by colonoscopy/FCP |
| Remission IBD | 19 | 2M, 4F, 13 not reported | 37-69 | n=10 Ulcerative colitis n=9 Crohn’s disease |
| Active IBD | 6 | 1M, 2F, 3 not reported | 37-43 | n=1 Ulcerative colitis n=5 Crohn’s disease |

**Supplementary Table 3:** Genes most significantly upregulated in AKR mice and downregulated in BALB/c mice as a result of 24 hours *T. muris* infection.

| **Gene name** | **Gene symbol** | **Log2 fold change** | **1-IPPLR** |
| --- | --- | --- | --- |
| Receptor for advanced glycation end-products | *Rage* | 2.071838311 | 2.57E-03 |
| Predicted gene 3579 | *Gm3579* | 1.742601729 | 3.89E-03 |
| Cadherin-related family member 4 | *Cdhr4* | 1.420705862 | 1.57E-03 |
| Anti-Mullerian hormone type 2 receptor | *Amhr2* | 1.384304979 | 1.03E-04 |
| Lactate dehydrogenase C | *Ldhc* | 1.322490739 | 5.86E-04 |
| V-myc myelocytomatosis viral related oncogene, neuroblastoma derived (avian) | *Mycn* | 1.322458717 | 1.89E-03 |
| Methyl-CpG binding domain protein 1 | *Mbd1* | 1.307428002 | 1.11E-02 |
| Phospholipase A2, group IIA (platelets, synovial fluid) | *Pla2g2a* | 1.301640648 | 3.14E-02 |
| GRP1 (general receptor for phosphoinositides 1)-associated scaffold protein | *Grasp* | 1.290711674 | 2.14E-02 |
| Apolipoprotein B mRNA editing enzyme, catalytic polypeptide 2 | *Apobec2* | 1.282355534 | 4.33E-02 |
| Fatty acid desaturase 2 | *Fads2* | 1.282163113 | 3.40E-04 |
| Cell division cycle 26 | *Cdc26* | 1.247747971 | 3.15E-03 |
| Karyopherin (importin) alpha 1 | *Kpn*α*1* | 1.205551397 | 5.11E-03 |
| Unknown | Unknown | 1.203247162 | 3.17E-02 |
| Tubulin tyrosine ligase-like family, member 10 | *Ttll10* | 1.15727629 | 3.98E-02 |
| Unknown | Unknown | 1.120285547 | 2.58E-02 |
| Intermediate filament family orphan 1 | *Iffo1* | 1.054095692 | 3.63E-02 |
| COP9 (constitutive photomorphogenic) homolog, subunit 3 (*Arabidopsis thaliana*) | *Cops3* | 0.982700593 | 6.91E-03 |
| Rhesus blood group-associated A glycoprotein | *Rhag* | 0.972268205 | 4.67E-02 |
| Uroplakin 1A | *Upk1a* | 0.946134944 | 3.23E-02 |

**Supplementary Table 4:** Genes most significantly downregulated in AKR mice and upregulated in BALB/c mice as a result of 24 hours *T. muris* infection.

| **Gene name** | **Gene symbol** | **Log2 fold change** | **IPPLR** |
| --- | --- | --- | --- |
| Haptoglobin | *Hp* | -2.629376068 | 5.50E-03 |
| Inhibitor of kappa B kinase beta | *Iκbkβ* | -2.618992274 | 2.23E-02 |
| Orosomucoid 1 | *Orm1* | -1.80638621 | 3.41E-02 |
| DNA segment, Chr 12, ERATO Doi 123, expressed | *D12Ertd123e* | -1.455622365 | 5.58E-05 |
| RIKEN cDNA A530053G22 gene | *A530053G22Rik* | -1.419325765 | 3.46E-02 |
| Intersectin 1 (SH3 domain protein 1A) | *Itsn1* | -1.318574188 | 4.48E-02 |
| Leptin | *Lep* | -1.210432562 | 1.89E-02 |
| Embigin | *Emb* | -1.144217553 | 1.74E-02 |
| Unknown | Unknown | -1.110624058 | 4.49E-03 |
| RAB14, member RAS oncogene family | *Rab14* | -1.084232232 | 6.42E-03 |
| Melanocortin 2 receptor accessory protein | *Mrap* | -1.05187992 | 1.36E-02 |
| Suppressor of cytokine signalling 7 | *Socs7* | -1.023987407 | 3.35E-02 |
| Establishment of cohesion 1 homolog 2 (*S. cerevisiae*) | *Esco2* | -0.989884374 | 3.76E-02 |
| Maestro heat-like repeat family member 2A | *Mroh2a* | -0.967874212 | 3.19E-02 |
| Solute carrier family 25 (mitochondrial carrier, Aralar), member 12 | *Slc25a12* | -0.934459822 | 9.07E-03 |
| Unknown | Unknown | -0.929840778 | 3.56E-02 |
| Prostaglandin E receptor 3 (subtype EP3) | *Ptger3* | -0.924710281 | 1.72E-02 |
| ADP-ribosyltransferase 3 | *Art3* | -0.914375161 | 4.73E-02 |
| Mesenchyme homeobox 2 | *Meox2* | -0.909832736 | 3.57E-02 |
| Prostaglandin E synthase | *Ptges* | -0.894574148 | 2.60E-02 |

13.	*MATLAB*. 2010, The MathWorks Inc.: Natick, Massachusetts.

**Supplementary figure legends**

**Supplementary Figure 1: Colitis-susceptible AKR mice show delayed DC response and increased eosinophilia in MLN after *Trichuris muris* infection**. (A-D) Dendritic cell (CD45^+^ MHCII^+^ CD11c^+^ F4/80^-^ CD103^+/-^ CD11b^+/-^), macrophage (CD45^+^ MHCII^+^ F4/80^+^ CD11c^+/-^), neutrophil (CD45^+^ Ly6G^+^ CD11b^+^ CD115^-^) and eosinophil (CD45^+^ Siglec-F^+^ CD11c^-^) populations as proportion of CD45^+^ cells (±SEM) in naïve mice and during *T. muris* challenge. n=3 mice per time point. Analysis by two-way ANOVA with Sidak’s multiple comparisons post hoc test. *P<0.05.

**Supplementary Figure 2: Scatter plot of changes in the expression of transcription factors calculated in TIGERi (TFA illustrator for global explanation of regulatory interactions).** Gene IDs in red indicate transcription factor upregulation and gene IDs in blue indicate transcription factor downregulation in AKR (A) and BALB/c (B) mice following 24-hour infection with Trichuris muris. n=4 mice per time point.

**Supplementary Figure 3: No difference in serum and faecal levels of calprotectin subunit S100A8 in colitis resistant or susceptible mice and poor correlation with colitis score.** Serum (A) and faecal (B) S100A8 levels measured by ELISA at 1, 7 and 21 days post infection (±SEM). Correlation of serum (C) and faecal (D) S100A8 versus colitis score at 0, 1 and 21 days post-infection. n=3-5 mice per time point. Analysis by linear regression, two-way ANOVA with Sidak’s post hoc test.

**Supplementary Figure 4:** **Membrane-bound ADAM10 expression in the proximal colon following *Trichuris muris* infection.** Representative images of proximal colon from naïve and *T. muris* infected AKR and BALB/c mice at 21 days post infection. Sections are stained for nuclei (DAPI; blue), epithelial cytokeratin (FITC; green) and ADAM10 (AF555; red). Inset area shows increased ADAM10 expression in the immune cells of the lamina propria. Bar=100μm, inset=50μm^2^.
