## Supplementary figures and images for "Differential expression of soluble receptor for advanced glycation end-products (sRAGE) in mice susceptible or resistant to chronic colitis"

### Supplementary Figure 1

#### Supplementary Figure 1

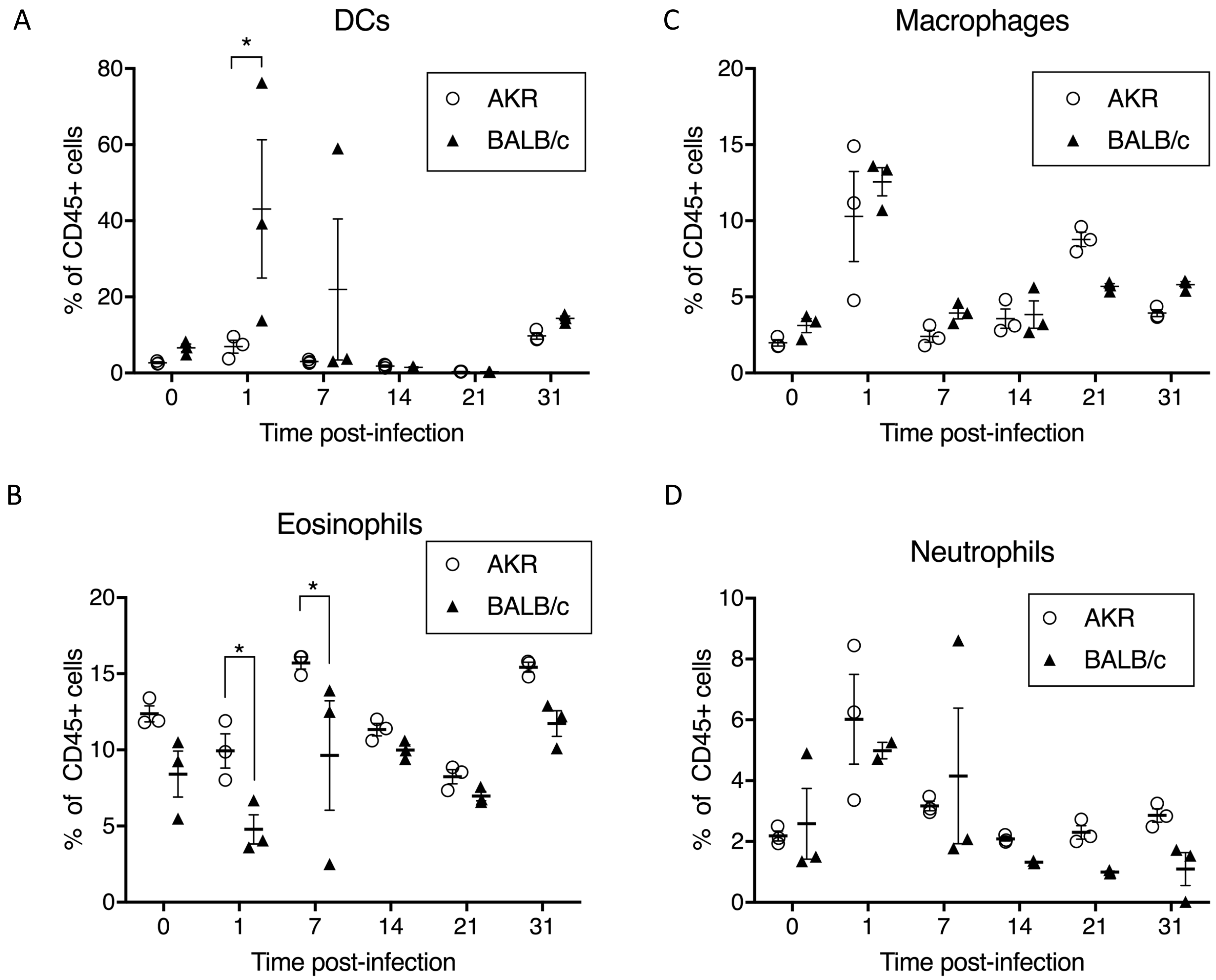

### Supplementary Figure 2

Supplementary Figure 2

A

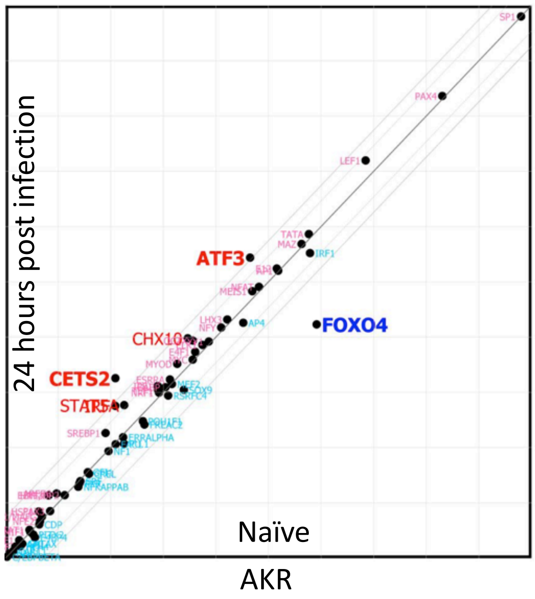


B

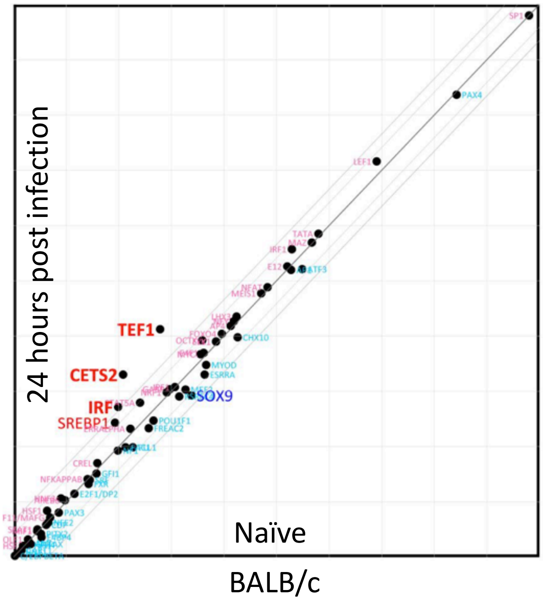

### Supplementary Figure 3

#### Supplementary Figure 3

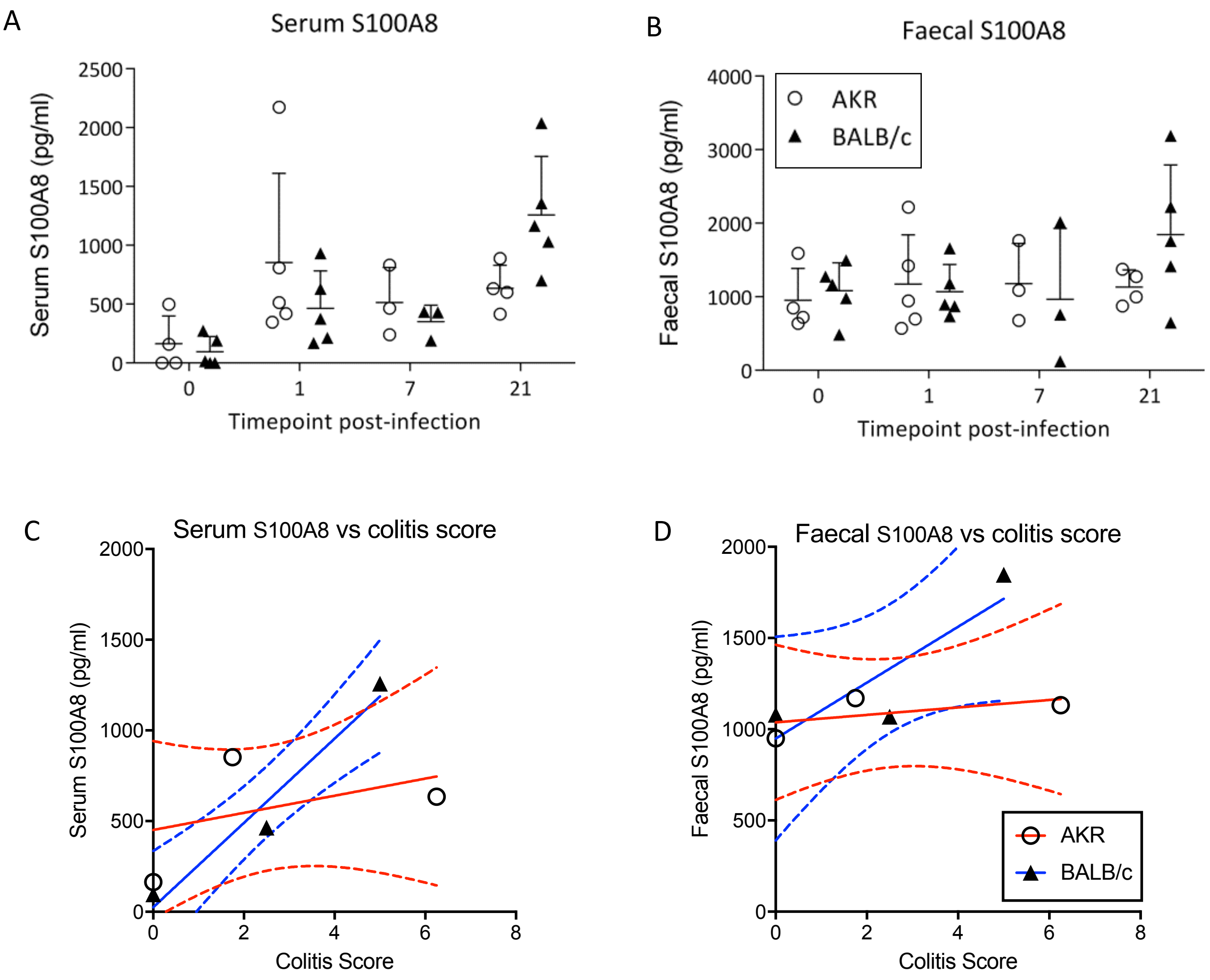

### Supplementary Figure 4

#### Supplementary Figure 4

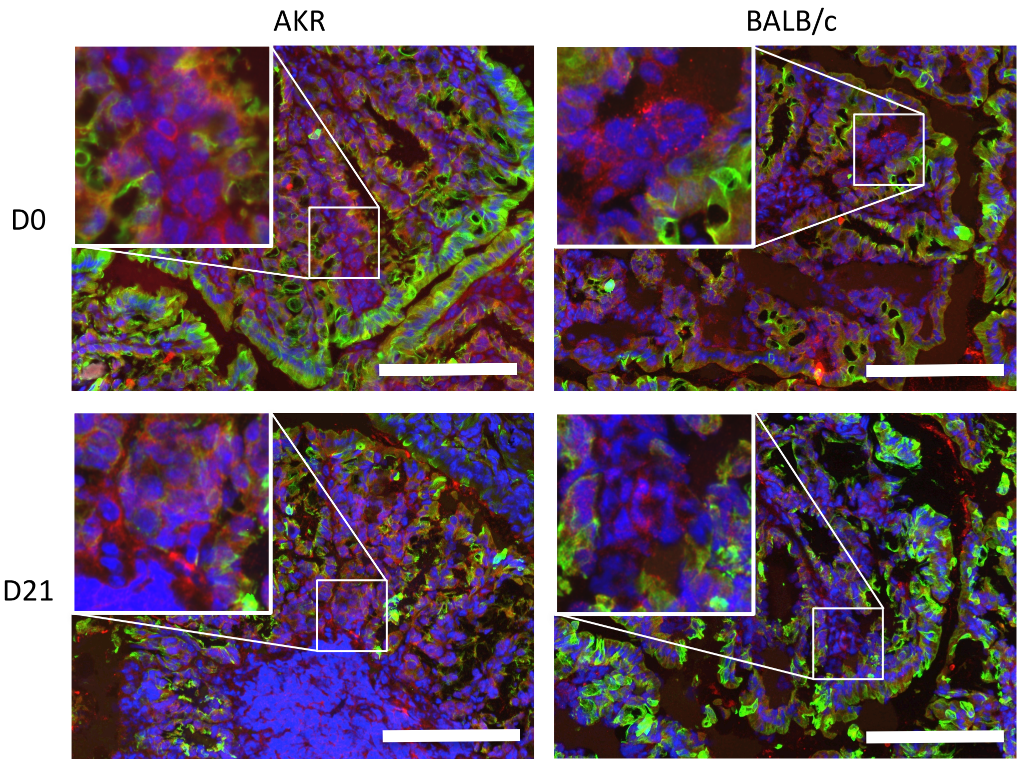
